# DNA StairLoop: Achieving High Error-correcting and Parallel-processing Capabilities in DNA-based Data Storage

**DOI:** 10.1101/2024.11.07.622581

**Authors:** Zihui Yan, Guanjin Qu, Xin Chen, Gang Zheng, Huaming Wu

## Abstract

DNA-based data storage is a promising solution to the challenges of large-scale data storage. However, the low throughput of the mainstream inkjet-based DNA synthesis method has hindered its widespread adoption. In contrast, high-throughput electrochemical synthesis provides higher throughput but with more nucleotide insertion, deletion, and substitution errors. Here, we propose an innovative coding scheme with high error correction capabilities, called DNA StairLoop. This coding scheme features a staircase interleaver and allows for component codes such as the convolutional code and the ow-density Parity-check code, allowing for flexible adaptation of the coding scheme. Both the row and the column decoders are soft input and soft output, enabling further improvement in data recovery accuracy through iterative decoding. The staircase interleaver facilitates extensive parallel decoding capabilities while effectively preserving parallelism across a multitude of nodes. In the in vitro experiment, DNA StairLoop successfully recovered the raw information in a staircase block with a nucleotide error rate of more than 8%. The simulations revealed that the DNA StairLoop can correct 10% nucleotide errors. Moreover, in parallel computing processing, the decoding time of our code continues to decrease dramatically.

## Introduction

In recent years, the unique properties of Deoxyribonucleic acid (DNA), including its environmental friendliness [1], long-term stability [2–4], and ability to store vast amounts of information in a compact form [5–8], have made it an attractive medium for data preservation. This has provided a promising solution to the growing demand for large-scale data storage. In this community, DNA synthesis, sequencing, and error-correction methods collaborate synergistically [9]. The use of error correction codes to correct nucleotide errors introduced during DNA synthesis and sequencing techniques has enabled reliable data recovery, rendering DNA-based data storage a feasible reality. This collaboration has further stimulated interest in the field [10].

However, the high cost associated with traditional inkjet-based DNA synthesis methods poses significant challenges to the widespread adoption of DNA-based data storage [11]. The inkjet printing-based DNA synthesis method requires complex equipment, which presents challenges for scaling up throughput. Compared to inkjet printing-based synthesis, which is expected to synthesize one million oligos per chip [12], high-throughput electrochemical synthesis based on micro semiconductor chip (MSC) technology has the potential to produce up to 200 billion oligonucleotides per chip at a cost as low as $50 per terabyte of data [13]. In 2021, Nguyen *et al*. developed a nanoscale DNA storage writer using an electrode array with a density of 25 × 10^6^ sequences per square centimeter [14]. High-throughput electrochemical DNA synthesis has thus emerged as a promising approach to increasing throughput and reducing costs. However, electrochemical synthesis is associated with a higher nucleotide error rate, which complicates the reliable storage and retrieval of data. The synthesized oligos exhibit approximately 8% insertion, deletion, and substitution (IDS) errors in the study of Nguyen *et al*. [14]. While this poses challenges for applications such as primer synthesis, in the context of DNA-based data storage, IDS errors can be mitigated through the use of error correction codes.

Recent advancements in DNA-based data storage have highlighted the need for robust error correction codes capable of correcting errors inherent in DNA synthesis, including nucleotide insertions, deletions, and substitutions. Traditional codes such as Reed-Solomon (RS) codes [8, 10, 11], Low-density Parity-check (LDPC) codes [15], and cyclic redundancy check (CRC) codes [16, 17], have been employed to detect errors. Moreover, several new IDS-specific error correction codes have been designed to correct IDS errors effectively. Notable examples include the Varshamov-Tenengolts (VT) code [18], the watermarker code [19], the HEDGES code [9], the DNA-Aeon code [16], and the Spider-web code [20]. The robustness of these codes has been validated in in vitro experiments, primarily via inkjet printing synthesis platforms. Additionally, although some codes have not yet been tested in wet experiments, simulations indicate that IDS-specific error correction codes, such as the single IDS error correction code [21], the multiple IDS error correction code based on number theory [22], and the time-varying code [23], possess strong error correction capabilities. However, the error-correction capabilities of the aforementioned codes are limited, as none can correct more than 8% of IDS errors, which aligns with the IDS error rates observed in our electrochemical synthesis experiments. An alternative approach to data recovery involves multiple synthesis attempts and increasing sequencing coverage to select the correct sequence from numerous sequencing reads [11, 24, 25]. Nevertheless, achieving correct sequence synthesis remains challenging for methods with high error rates. For example, in a storage experiment conducted by Antkowiak *et al*. [11], only one out of three files was successfully recovered, even with high sequencing coverage.

The majority of the aforementioned coding schemes utilize concatenated codes in the form of block interleavers, with the outer code correcting dropout errors and the inner code correcting or detecting IDS errors. However, block interleavers inherently lack support for parallel decoding, which constrains improvements in data retrieval speed. To facilitate parallel decoding, the data block must be subdivided into smaller, independent blocks, allowing concurrent decoding of each block. In this approach, however, the encoding and decoding processes of individual sub-blocks are independent, precluding the exchange of information between them that could otherwise enhance correction capabilities. Furthermore, studies have shown that the accuracy of DNA synthesis and sequencing is closely linked to the biochemical properties of oligos, such as high GC content, long homopolymers, and the presence of undesired motifs [26–28]. Designing a coding scheme that simultaneously meets biochemical constraints while enabling parallel decoding and achieving high error-correction performance is a significant challenge for DNA-based data storage.

In this study, we propose a novel encoding scheme for DNA-based data storage, called DNA StairLoop, which offers high error-correcting capability and supports highly parallel decoding. Specifically, this scheme is optimized for high-throughput electrochemical DNA synthesis through the following features: (i). The encoding structure utilizes a staircase interleaver [29], enabling information exchange between data blocks to enhance overall error resilience. The row and column component codes can differ, allowing the flexible use of various error-correcting codes such as convolutional codes and LDPC codes. (ii). A soft-input soft-output (SISO) row decoder is employed to correct IDS errors in both convolutional and LDPC codes, supporting iterative decoding to improve error-correcting performance. (iii). The staircase interleaver enables highly parallel decoding. In our experiments, this parallel decoding capability, based on the interleaver’s structure, resulted in significant time reduction, even when tested across thousands of nodes, demonstrating its potential to effectively reduce decoding time. Additionally, we propose an extended encoding scheme using the convolutional code with a code rate of 1*/*3, which maintains a GC content between 33.3% and 66.6% within a sliding window and prevents the formation of homopolymers exceeding three consecutive nucleotides. In our experiments, we adopted it as the row code and the LDPC code as the column code. The effectiveness and robustness of our coding scheme have been verified by in silico and in vitro storage experiments. Simulations showed that our scheme can achieve a high IDS error tolerance, enabling data recovery when the IDS error rate is 10%. In in vitro storage experiments, the DNA StairLoop accurately recovers the original information from a block with an IDS error rate of 8.13%.

## Results

### Coding Structure Overview

The high-throughput electrochemical synthesis approach has a significantly higher throughput threshold, exceeding that of inkjet printing-based by five orders of magnitude (Fig. 1a). However, its nucleotide IDS error rate is also relatively high. To address this issue, we propose an innovative DNA coding scheme, DNA StairLoop. It has five important features:

1) The original message is written in a staircase interleaver, in which the connections between successive data bit matrices are of the staircase type (Fig. 1b).
2) The encoding structure is a serial-concatenated code with independent row codes and column codes. The component codes can be convolutional codes or block codes. For different combinations of component codes, the encoding and decoding schedules are distinct, as shown in Section 1 of the Supplementary Materials (Fig. S1).
3) The decoder is iterative and follows the turbo principle. Specifically, the row decoder and column decoder are both soft-in and soft-out, which iteratively feed the probabilities of the information bits back and forth to each other (Fig. 1c).
4) The row decoder is specifically designed for DNA-based data storage. It can work in conjunction with the BCJR algorithm for convolutional codes and the sum-product algorithm for LDPC codes, enabling the traditional decoding algorithms to correct insertion and deletion (indel) errors (Fig. 1d).
5) The staircase matrix divides the parity nodes into parallel nodes. The nodes decode and pass outer-matrix information from the first and last ends to the upper and lower nodes. The DNA StairLoop scales well in parallel because the information transfer involves only a single matrix and uses nonblocking communication (Fig. 1e).

**Fig 1.**
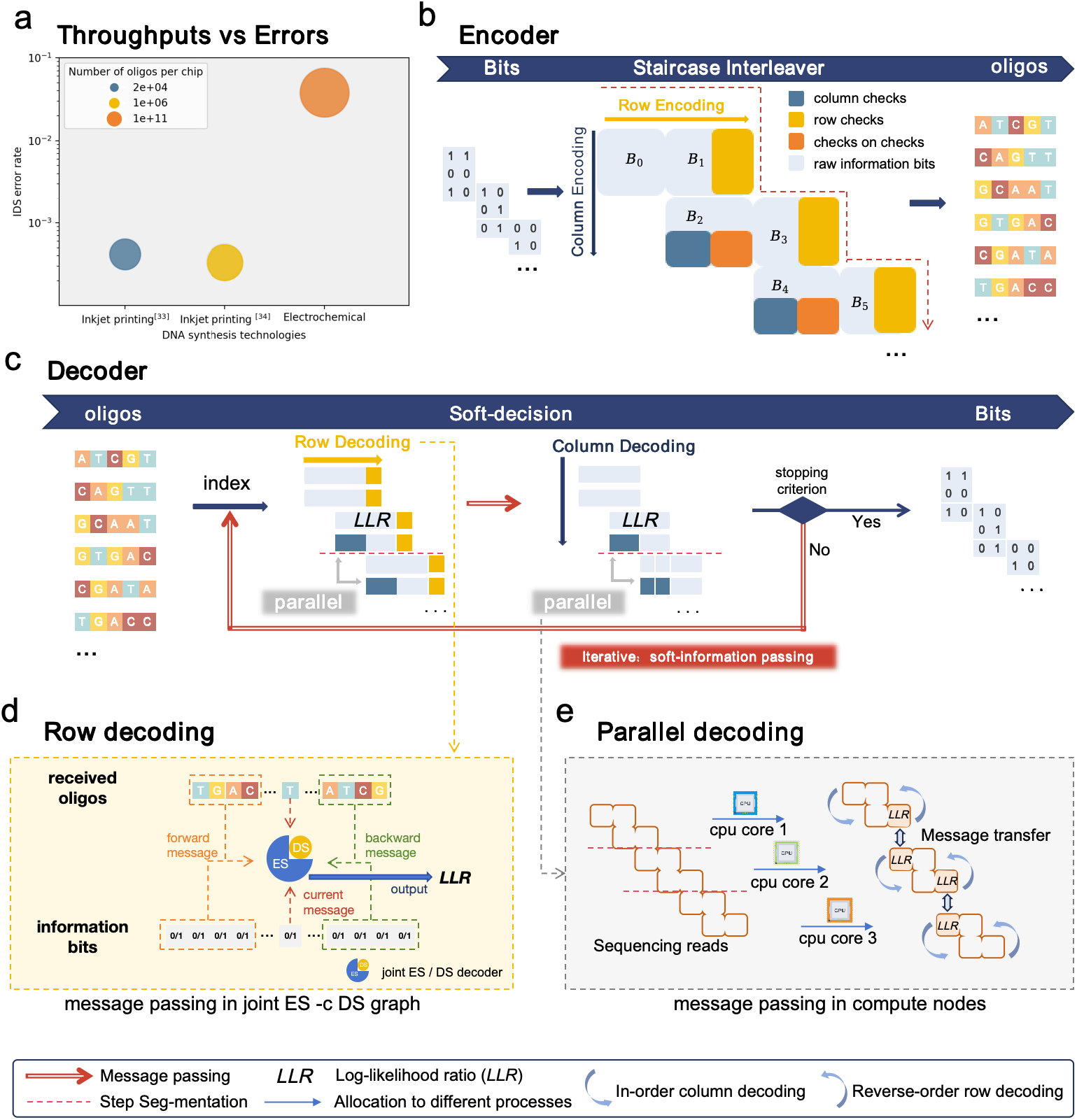
Overview of the DNA StairLoop coding scheme. (a).Distribution of nucleotide IDS error rates and throughput for inkjet printing-based synthesis by Agilent [30] and Twist [12], and electrochemical synthesis by GenScript [13]. **(b)**. Illustration of the encoding structure. The raw binary data are first divided into short sequences and then sequentially input into the blocks (in the positions of the raw information bits), forming a staircase layout. The raw information blocks subsequently undergo row encoding and column encoding to generate codewords, i.e., oligos. **(c)**. Diagram of the decoding structure. The sequencing data first undergo raw decoding and are subsequently restored to the staircase blocks according to their indices. The row decoder is soft-input and soft-output, producing extrinsic information in the form of the log-likelihood ratio (LLR) of the raw information bits. The column decoder receives this extrinsic information and performs soft-decision column decoding, which outputs extrinsic information in the form of the LLR of the raw information bits as well. This extrinsic information is iteratively input back into the row decoder. The message passing path follows the red arrows shown in the diagram. The iteration between the row and column decoders continues until the stopping criterion is met, ultimately producing the decoded bits. **(d)**. Trellis diagram of the row decoding scheme. The forwards message, backwards message, and current message are obtained from the trellis by tracing all possible trellis paths in the joint encoding state (ES) - drift state (DS) graph. The LLR of the *i*-th information bit is subsequently calculated on the basis of the forwards message, backwards message, and current message in the *i*-th decoding moment. **(e)**. Flowchart for parallel decoding. The stepped matrix is equally distributed to different nodes. Row and column decoding is performed within each node, and the first and last outer information matrix that is decoded is passed to the previous and next nodes.

### Encoding Scheme and Staircase Interleaver

The staircase interleaver is a generalized block type interleaver. The raw binary data are written into the information matrices in a predefined order *B*_1_, *B*_2_, …, *B*_*F* −1_, and arranged in a staircase pattern, as shown in Fig. 1b. According to the structure of the staircase interleaver, the encoding process is performed in two phases, i.e., row encoding and column encoding. For convenience, we call the column encoder “*E*_*c*_” and the row encoder “*E*_*r*_”. Herein, *E*_*r*_ and *E*_*c*_ can be chosen from convolutional codes and block codes (especially the LDPC codes) and can even be different. We provided four interleavers with different component codes, as shown in Section 1 of the Supplementary Materials. For different component codes, the arrangement of information bits varies significantly depending on the design of the interleaver. This is because, in the iterative decoding process, the block codes require extrinsic information for all the codeword bits, whereas the convolutional codes require only systematic information bits. This also causes the encoding schedules to be different. However, transmitted codewords are fixed as row sequences.

Encoding proceeds recursively on each matrix pair. First, matrices *B*_0_ and *B*_*F*_ are initialized to reference states known to the encoder and decoder, e.g., matrices of all zero symbols. Two error-correction codes in the systematic form are given: the row code and the column code. For *i* = 1, …, ⌈(*F* − 1)*/*2⌉, row encoding proceeds on the matrix pair [*B*_2*i*−2_ *B*_2*i*−1_], generating the row codeword matrix pair 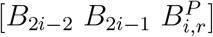. The additional elements in the *j*-th row of 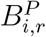 are the parity symbols that result from encoding the information symbols in the *j*-th row of [*B*_2*i*−2_ *B*_2*i*−1_]. Column encoding proceeds on the matrix pair [*B* _2*i*−1_ *B*_2*i*_]^*T*^, generating the column codeword matrix pair 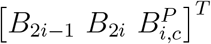. The additional elements in the *j*-th column of 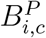 are the parity symbols that result from encoding the information symbols in the *j*-th column of [*B*_2*i*−1_ *B*_2*i*_]^*T*^. Herein, the relationship between successive matrices satisfies the following relation: each of the rows is a valid codeword of the row encoder, and each of the columns is a valid codeword of the column encoder.

According to the relationships of the codewords, the connections within the staircase interleaver are clearly defined. We call two consecutive matrices of the same layer as a block, such as [*B*_0_, *B*_1_]. On the one hand, the parity-check information is transmitted between the staircase blocks, allowing the encoder to emulate the performance of long codes. On the other hand, the independence of nonadjacent blocks within the staircase interleaver enhances the parallelism of the decoding process.

### Iterative Decoding Scheme

The key behind DNA StairLoop is the soft-decision decoders, which help improve the error correction capabilities through iterative decoding. Like the encoder, we call the column decoder “*D*_*c*_” and the row decoder “*D*_*r*_”. Since DNA strands suffer IDS errors in transmission, the row decoder *D*_*r*_ aims to regain synchronization or to correct IDS errors. The column decoder *D*_*c*_ uses the soft output of the row decoder to compute likelihood information without suffering from IDS errors. In this case, we propose an IDS error correction strategy to improve the row decoder, which is presented in detail below. However, the BCJR algorithm for the convolutional code and the sum-product algorithm for the LDPC code can be easily applied to the column decoder with a few minor adaptations. Given that our coding scheme is a generalized serial connection and that *D*_*c*_ does not directly observe the transmission information of the channel, it relies on the information received from *D*_*r*_ to output the log-likelihood ratio (LLR) on the *E*_*c*_ output bits. *D*_*r*_ receives the channel LLR on the received bits and the extrinsic information from *D*_*c*_ to calculate a new LLR value.

Consider the row decoder *D*_*r*_ as an example. Let ***v*** = {*v*_1_, *v*_2_, …, *v*_*n*_} ∈ {0, 1}^*n*^ and ***r*** = {*r*_1_, *r*_2_, …, *r*_*N*_ } ∈ {0, 1}^*N*^ denote the channel inputs of length n and outputs of length N, respectively. Let *S* denote the encoding syndrome constraint. Then the posterior L-value of *v*_*t*_ can be written as

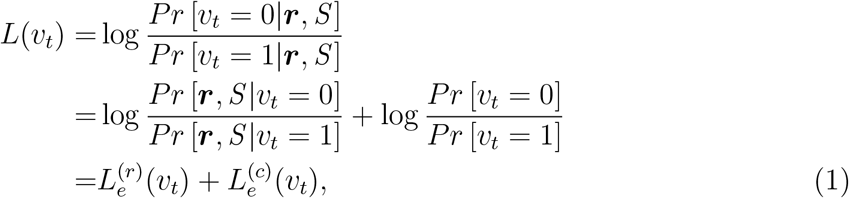

of which only 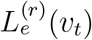 is passed to *D*, and 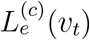 is the interleaved extrinsic information from the previous *D*_*c*_ iteration.

The outline of the iterative decoding scheme is as follows:

1) All the metrics are initialized appropriately, and the extrinsic information is set to zero.
2) Row decoder: The row decoding algorithm is run with received bits and the extrinsic information from *D*_*c*_ to obtain the soft decision of the information symbols of *E*_*r*_, and the extrinsic information is sent to *D*_*c*_.
3) Column decoder: The column decoding algorithm is run with the extrinsic information from *D*_*r*_ to obtain the soft decision of the output symbols of *E*_*c*_, and the extrinsic information is sent to *D*_*r*_.
4) Steps 2 and 3 are repeated until the preset maximum number of iterations is reached (or the stopping criterion is satisfied). Decisions are made on information bits according to the final soft decision computed by *D*_*c*_.

### Row decoding strategy

The processes of DNA synthesis and sequencing introduce nucleotide IDS errors within a single DNA strand, which are typically modeled as an IDS channel (Fig. S2) [31, 32]. The input to the row decoder is a DNA sequencing read containing IDS errors. Consequently, the row decoder must be capable of correcting IDS errors and re-establishing sequence synchronization.

In this work, we provide an IDS error-correction strategy to modify the conventional decoding algorithms of the convolutional code and the LDPC code to correct IDS errors. The IDS channel model can be conveniently represented by a trellis, where each input *v*_*i*_ corresponds to a trellis node (Fig. S3). However, unlike the discrete-time intersymbol interference channel [33], each trellis node of the IDS channel model cannot correspond to a single output because of indel errors. To cope with indel errors, we introduce a new hidden state variable into the trellis graph, called the synchronization drift. For *t* = 1, 2, …, *n*, the drift state is defined as 𝒟_*t*_ at time *t*, and its realization *d*_*t*_ is defined as the number of insertions minus the number of deletions that occurred until symbol *v*_*t*_ is transmitted. In addition, we define the encoding state at time *t* as 𝒮_*t*_, of which the realization *s*_*t*_ is determined by the encoding scheme.

The key innovation behind the row decoding strategy is the transmission of the encoding state (ES) and the drift state (DS), which facilitates the maximum a posteriori (MAP) decoding. Consider the MAP decoding criterion, which is given by

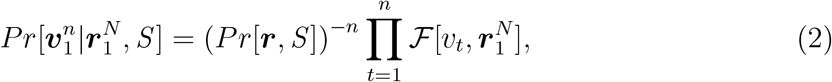

where each factor is

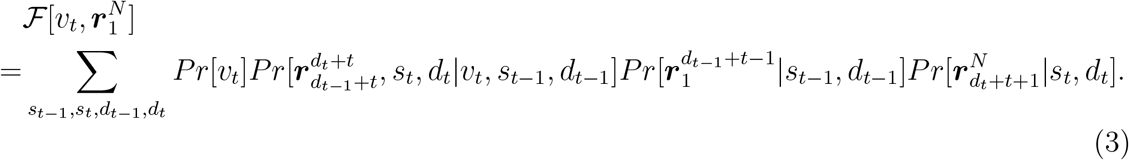

Each factorization can be illustrated in the factor graph (Fig. S3), where for each factor of Eq. (3), there is a single trellis node, and for each variable on which the factor depends, there is a single connected edge. Armed with the above graph representation of the trellis, the message-passing algorithms can be used on IDS channel models. Then, the maximum posterior probability of each input symbol can be calculated. The details of the decoding strategy are presented in Section 2 of the Supplementary Materials.

In addition, by jointly calculating the maximum APP of each input symbol on multiple copies, the decoder can obtain higher performance gains from the sequencing redundancy and skip the complex multiple sequence alignment (MSA) process. In contrast to MSA technology, which makes hard decisions on the basis of majority voting [34, 35], this decoding strategy outputs soft information, which is more applicable to the iterative decoding scheme.

### Parallel decoding scheme

The DNA StairLoop adopts a parallel decoding method that acts on the staircase interleaver, which enables the parallel decoding of thousands of nodes to increase the decoding speed. The decoding process of the DNA StairLoop is divided into two parts: local decoding and communication. When decoding, the ladder blocks are evenly divided and distributed to parallel nodes, and each parallel node performs inverse-order row decoding and sequential column decoding in its local decoding. When a node finishes decoding the last set of blocks in the inwards row, it passes the outer information to the next node and continues to decode the previous blocks in the inwards row, which does not hinder the decoding process because the nonblocking time passes. Column decoding also passes the outer information to the previous node after decoding the first block and continues with column decoding. We experimentally verified that parallelism can be maintained, even with thousands of nodes. This helps reduce the time-consuming decoding problem in large-scale storage.

### Performance analysis of the DNA StairLoop with simulated data

We performed simulations to evaluate the effectiveness of the DNA StairLoop and compared its error correction performance with that of state-of-the-art algorithms such as DNA-Aeon and DNA Fountain (Table 1). We first evaluated the error-correction capabilities of the DNA Fountain code, DNA-Aeon code, and our code under varying error rates (Fig. 2a). To ensure a fair comparison, we adjusted the encoding parameters so that all codes operated at nearly identical code rates. The details of the experimental setup are shown in Section 4.1 of the Supplementary Materials. The IDS error rates used in the simulations are consistent with those observed in our electrochemical synthesis experiment. The DNA StairLoop can successfully decode information with an error rate as high as 10%, and its maximum error resilience exceeds that of DNA-Aeon and DNA Fountain. These findings demonstrate that our code is a viable solution for error-prone synthesis methods. In addition, we verified in the Supplementary Materials that the error-correction capability of the DNA StairLoop degrades when the staircase interleaver is not used, which proves that the staircase interleaver can improve the error-correction capability. Iterative decoding is a key factor in the high error-correction capability, and we show its effect with increasing error rates in Fig. 2c. As the number of iterations increases, the decoding recovery rate continues to improve. Even for an unpredictable high error rate, the decoded recovery rate can still be improved through iteration. For example, when the IDS error rate is 11%, the decoding effect is significantly improved as the number of iterations increases. In addition, the direct input of DNA StairLoop can be duplicated sequencing reads rather than the central sequence obtained by clustering methods. Fig. 2d shows the effect of the copy number on the decoded recovery rate, with an IDS error rate of 11%.

**Table 1.** Comparison among popular DNA coding schemes and key achievements.

|  | DNA Fountain [10]<br>(Erlich and Zielinski) | HEDGES [9]<br>(Press et al.) | DNA-Aeon [16]<br>(Welzel et al.) | DNA StairLoop<br>(This work) |
| --- | --- | --- | --- | --- |
| Coding scheme<sup>1</sup> | inner: RS code | inner: HEDGES | inner: AC based | **row: extended convolutional code** |
|  | outer: fountain code | outer: RS code | outer: Raptor code | **column: LDPC** |
| Interleaver<sup>1</sup> | block type | block type | block type | **staircase type** |
| Constraints<sup>1</sup> | homopolymers | homopolymers | homopolymers | **homopolymers** |
|  | GC | GC | GC, motifs | **GC, motifs** |
| Synthesis methods<sup>1</sup> | inkjet printing-based | inkjet printing-based | inkjet printing-based | **high-throughput electrochemical** |
| Synthesis throughput<br>(distinct oligos per chip)<sup>2</sup> | $\leq 696,000$ | N/A | N/A | **8,000,000** |
| Error-correction<br>capability<sup>3</sup> | 2% | N/A | 8% | **10%** |
| Parallel decoding<sup>1</sup> | N/A | N/A | local parallel<br>(inner decoder) | **global parallel**<br>**(inner+outer decoder)** |

<sup>1</sup> The information presented is derived from the original data provided in the references. These references employed different original designs, including input files, parameter designs, and experimental setups. This table is intended to provide a general comparative overview. The notation “N/A” indicates that the corresponding result is not available in the references.
<sup>2</sup> The throughput of DNA Fountain is from the calculation of Antkowiak *et al.* [11].
<sup>3</sup> The error-correction capabilities were evaluated through simulations, with the results obtained under nearly identical code rates across all codes.

**Fig 2.**
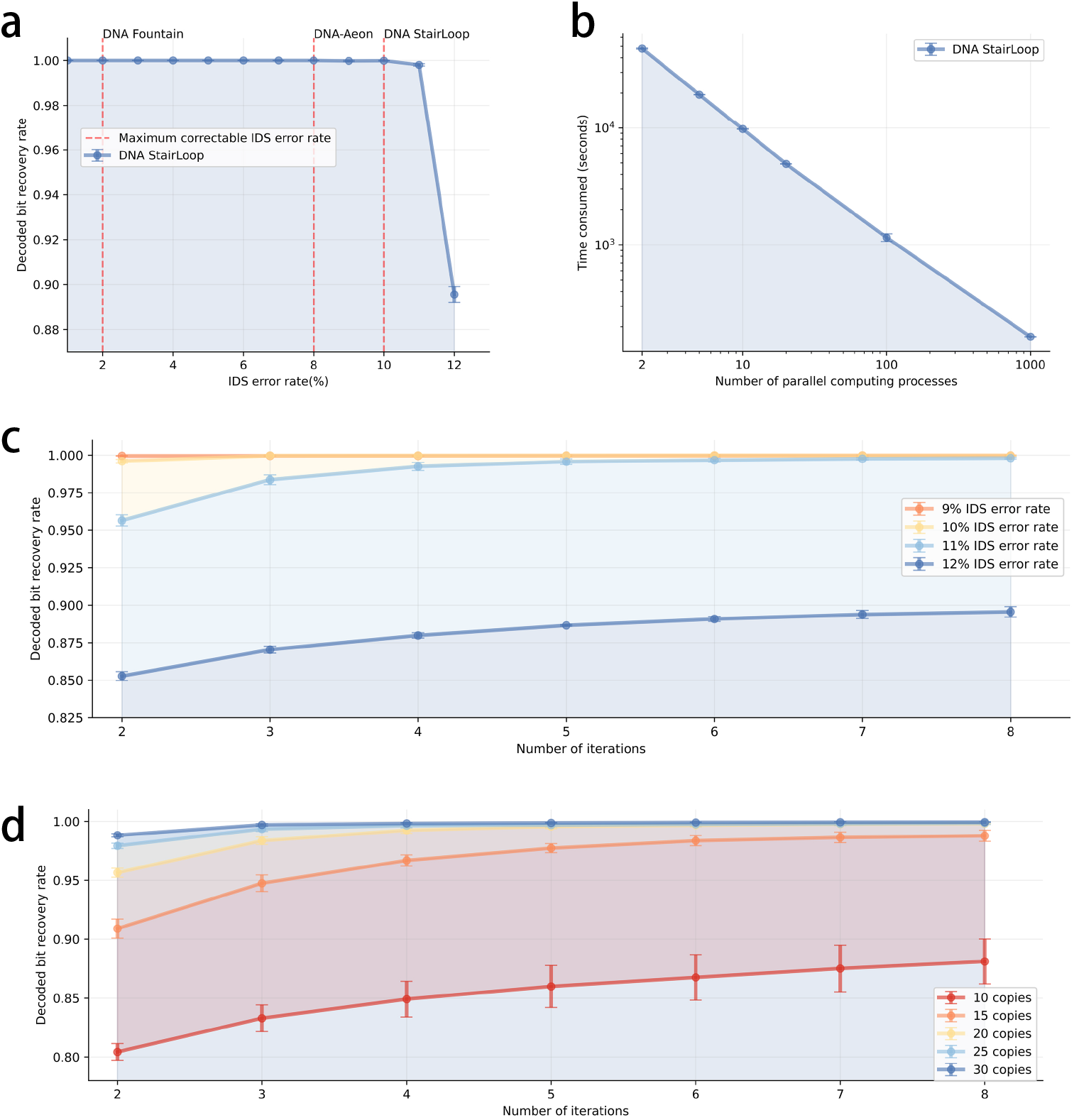
Simulation experiment results. (a).Percentage of Successful Decoding Attempts at a Given IDS Error Ratio. We simulate the recovery when storing 10 KB of data at 10 copy numbers, where the ratio of substitution, deletion, and insertion error rates is 0.25 : 0.61 : 0.12. For a fair comparison, the DNA Fountain code and DNA-Aeon code were configured to have similar code rates and GC content constraints as the DNA StairLoop. **(b)**. Impact of Increasing Parallel Nodes on Decoding Time. This parallel simulation was conducted on a supercomputer, with the number of blocks for DNA StairLoop set to 4000. **(c)**. Impact of Iterative Decoding on Recovery Rate. **(d)**. Impact of Copy Number on Recovery Rate. At this point, the substitution, deletion, and insertion error rates are 2.83%, 6.81%, and 1.35%, respectively.

As the amount of stored data increases, the decoding speed also becomes a critical concern. To address this, we tested the parallel decoding capabilities of DNA StairLoop. Fig. 2b illustrates the trend in decoding time as the number of nodes increases from 2 to 1,000. The results indicate a linear decrease in decoding time with the addition of nodes, demonstrating that DNA StairLoop maintains strong parallel efficiency, even at the scale of thousands of nodes. This scalability significantly reduces the overall decoding time.

Simulation experiments demonstrated that DNA StairLoop, which uses both iterative and multiple sequence decoding algorithms, exhibits exceptionally high error-correction capability, effectively addressing the challenges posed by high-error-rate synthesis methods. Moreover, its highly parallel decoding architecture substantially increases decoding speed, thereby minimizing the additional time costs typically associated with time-consuming decoding processes.

### Performance analysis of the in vitro experiments

We performed an in vitro storage experiment to verify the effectiveness of DNA StairLoop. In this experiment, three images and a text file were compressed to 740 KB and encoded into 14,810 sequences, each of which was 127 nucleotides in length, via an high-throughput electrochemical synthesis platform (Fig. S6). The encoding process utilized our extended convolutional code as the row code and a (100, 64) irregular LDPC code as the column code. Here, the short length of the LDPC code was deliberately chosen to increase the number of staircase blocks, enabling us to test the parallel computing capabilities of the system. Fig. 3 shows the detailed results of this in vitro experiment. The statistics of the sequencing reads indicate that the sequence length distribution under electrochemical synthesis conditions is more dispersed, with a significantly higher error rate than inkjet printing-based synthesis technology, which typically results in error rates well below 1%. This higher error rate was particularly pronounced for deletion and substitution errors (Fig. 3a and Fig. 3b). Further details on the experimental procedure and statistical methods are provided in Section 5 of the Supplementary Materials.

**Fig 3.**
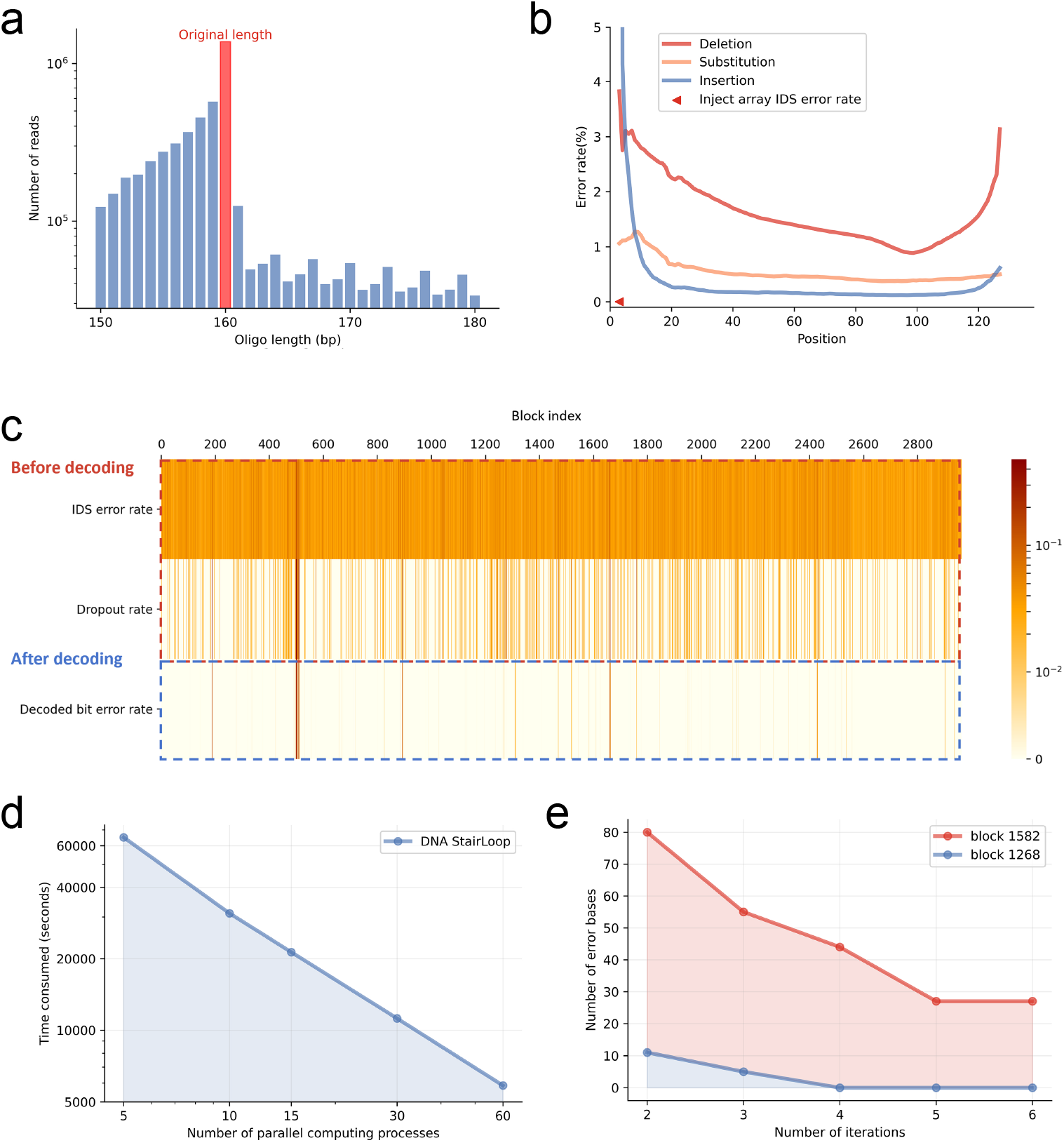
Results of the in vitro storage experiments. (a).Distribution of the lengths of sequenced sequences after double-ended splicing and filtering, with the original length set at 160 base pairs. **(b)**. Error rates at different positions within the sequence. **(c)**. Error rates, dropout rates, and final decoding accuracy for different blocks. A total of 740 KB of data was stored, divided into 2,962 blocks. Despite generally high error rates and some blocks with very high dropout rates, DNA StairLoop achieved a 99.8% recovery rate. **(d)** Variation in decoding time with the number of parallel processes. **(e)** Changes in the number of error bases with iterative decoding.

The block-by-block statistics of IDS error rates and decoded bit error rates demonstrate that DNA StairLoop can achieve robust decoding even with error rates exceeding 8%. For example, in the 93^*rd*^ block, with an error rate of 8.13%, a coverage rate of 7.66x, and a dropout rate of 2%, this block achieves error-free data recovery. This finding indicates that the DNA StairLoop possesses significant IDS error correction capability. However, owing to unforeseen circumstances, the dropout sequences exhibited significant clustering in blocks such as the 503^*rd*^ and 1661^*rd*^ blocks, where the sequence dropout rate exceeded 40%. This dropout rate is beyond the recovery capability of the (100, 64) LDPC code. Nevertheless, increasing the length of the LDPC code to prevent dropout sequences from clustering within a single block effectively mitigates these errors. We selected (600 and 1200) LDPC codes for the simulations, demonstrating that the parameter-adjusted DNA StairLoop can achieve error-free decoding under the high dropout rates observed in the experiments (see Section 4.2 of the Supplementary Materials). To test the parallelism of the decoding algorithm, we used a shorter LDPC code. Fig. 3d shows the correlation between parallel nodes and time spent, and shows that the decoding time can be significantly reduced with the increasing number of parallel nodes. In addition, taking two blocks as an example, the effect of iterative decoding is significant (Fig. 3e). As the number of iterations increases, the number of erroneous bases decreases significantly, and even for blocks that cannot be fully recovered due to high error and dropout rates, the DNA StairLoop can still reduce the number of erroneous bases as much as possible through iterative decoding. In conclusion, in the face of the high error rate and high concentration dropout rate caused by the current high-throughput electrochemical synthesis technology, the DNA StairLoop can still recover most of the raw information at a low decoding coverage rate (8.7x). For the whole file, the recovery rate is 99.8%. If the length of the column encoder is increased, the high concentration dropout rate can be further overcome, which proves that the DNA StairLoop is expected to be a potential encoding scheme for DNA-based data storage under high-throughput electrochemical synthesis conditions.

## Discussion

DNA-based data storage systems record information by writing it into a sequence of nucleic acid strands. Unlike conventional storage media, the synthesis and sequencing of nucleotide sequences leads to difficulty in writing and reading information, resulting in complex burst errors. This leads to challenges in the recovery of DNA data. The highest throughput electrochemical synthesis techniques currently available are also reported to produce many synthesis errors, notably with 6.3% deletion errors in nanoscale electrode synthesis [14] and 8.1% IDS errors in our in vitro experiments.

To achieve reliable information storage at such a high IDS error rate, we propose a staircase interleaver-based information encoding scheme and its soft-decision iterative decoding scheme. The use of the staircase interleaver ensures that the data matrices are both independent and transitive, making parallel computations with a low-error floor possible. On the decoding side, our proposed error correction strategy for the row component code can correct IDS errors. The iterative exchange of LLRs between row and column component codes can further enhance the error correction performance.

Notably, our proposed row decoding scheme can cooperate with traditional decoding algorithms for convolutional codes and block codes, enabling these codes, which can correct only substitution errors and erasure errors, to correct IDS errors. This makes it feasible to use convolutional codes or block codes as row codes. We propose four types of encoding structures for different row and column component codes. In the experiments, we employed an encoder using convolutional codes as row codes and LDPC codes as column codes. This is because encoders with convolutional codes acting as row codes can effectively implement artificially defined biochemical constraints, which is more challenging for LDPC codes. Furthermore, preliminary tests of the four encoding structures revealed that the combination of convolutional codes and LDPC codes has a greater error correction potential.

We performed in silico and in vitro experiments to test the error correction performance of our coding scheme. Compared with existing state-of-the-art codes, the DNA StairLoop has a significant advantage in terms of a high IDS error rate. If an unexpectedly high error rate is observed during decoding, the number of iterative decoding executions can be adjusted to improve the decoding performance. Multiple parameters of the codec, such as the number of blocks of the information matrix, the encoding structure of the row codes, the code rate of the column codes, and the number of iterative decodes, can be adjusted to further customize the base sequence constraints and improve the error correction capability, in accordance with the synthesis, storage, and sequencing methodology in use, as well as the expected decoding byte error rate requirements. In addition, if high IDS error rates are not taken into account during encoding, the number of iterative decoding executions can be adjusted to improve decoding performance.

In future work, we will address the challenge of continuous sequence dropout during DNA synthesis by extending the length of LDPC codes. Our simulations, as detailed in Section 4 of the Supplementary Materials, demonstrated that increasing the length of LDPC codes can eliminate the decoded byte errors caused by the high dropout rates observed in the in vitro experiments. Furthermore, we plan to investigate the integration of IDS error correction strategies with other error-correcting codes, such as Reed-Solomon (RS) codes, within the DNA StairLoop framework. We anticipate that the DNA StairLoop will significantly advance high-error-rate DNA synthesis technologies for DNA-based data storage.

## Data availability

The data generated in this work have been deposited in the Figshare database with the following DOI link:

The raw information stored in our in vitro experiment (https://doi.org/10.6084/m9.figshare.26212682.v1). The encoded oligos (https://doi.org/10.6084/m9.figshare.26212682). The raw sequencing reads (https://doi.org/10.6084/m9.figshare.26212682.v1). The cleaned sequencing data is used as input to the decoder (https://doi.org/10.6084/m9.figshare.26212718).

## Code availability

The code used by the DNA StairLoop can be found at https://github.com/Guanjinqu/StairLoop.

## Acknowledgements

This work was supported by the National Key Research and Development Program of China (no. 2020YFA0712100) and the National Natural Science Foundation of China (no. 62071327).

## Author contributions statement

H. W. and X. C. conceived and supervised the project. Y. Z. designed the encoding and decoding schemes. G. Q. derived analytical results and performed numerical calculations. G. Q. and Y. Z. analyzed the data. G. Q. and Y. Z. wrote the original draft, whereas H. W., X. C. and G. Z. reviewed and edited it. All authors read and approved the final paper.

